# From phages to mammalian viruses: viral receptors play a central role in protein-protein interaction network

**DOI:** 10.1101/727024

**Authors:** Fen Yu, Zheng Zhang, Yuanqiang Zou, Ye Qiu, Aiping Wu, Taijiao Jiang, Yousong Peng

**Affiliations:** College of Biology, Hunan University, Changsha, China; Changsha Qiangze Biotech Co., Ltd, Changsha, China; Suzhou Institute of Systems Medicine, Suzhou, China; Center of System Medicine, Institute of Basic Medical Sciences, Chinese Academy of Medical Sciences & Peking Union Medical College, Beijing, China

## Abstract

**Motivation:** Receptors on host cells play a critical role in viral infection. How phages select receptors is still unknown.

**Results:** Here, we manually curated a high-quality database named phageReceptor, including 355 pairs of phage-host receptor interactions, 280 unique viral species or sub-species and 64 bacterial species. Sugars and proteins were most widely used by phages as receptors. The receptor usage of phages in Gram-positive bacteria was different from that in Gram-negative bacteria. Most protein receptors were located on the outer membrane. The protein receptors were highly diverse in their structures, and had little homology with mammalian virus receptors. Further functional characterization of phage protein receptors in *Escherichia coli* showed that they had larger node degrees and betweennesses in the protein-protein interaction (PPI) network, and higher expression levels, than other outer membrane proteins, plasma membrane proteins, or other intracellular proteins. These findings were consistent with what observed for mammalian virus receptors, suggesting that viral protein receptors play a central role in the host’s PPI network. The study deepens our understanding of virus-host interactions.

**Availability:** The database of phageReceptor is publicly accessible at http://www.computationalbiology.cn/viralRecepetor/index.html.

## Introduction

Bacteriophages (phages) are a group of viruses that specifically infect bacteria or archaea ^[1, 2]^. Phages are the most abundant entities on the earth with diverse morphology, genomes, host range, and life cycles ^[3, 4]^. They play a key role in shaping bacterial population dynamics and balancing the global ecosystem ^[5, 6]^. For humans, phages, on one hand, cause large economic loss by killing engineered bacteria used in industries, such as the *Lactococcus lactis ^[7]^*; on the other hand, they can be applied for therapy of bacterial infections, especially for the super bacteria which are resistant to most antibiotics ^[8, 9]^. Besides, phages were also reported to impact human health ^[10, 11]^.

Attachment of phages to their specific receptors on host cells is the first step of phage infection ^[12, 13]^. The phage-receptor interaction is highly specific and dynamic ^[12, 14]^, and determines the host specificity of phages to a large extent ^[12]^. Receptors have been identified for lots of phages ^[13]^. Various biomolecules can be utilized by phages as their receptors, such as carbohydrate, lipid and protein ^[13, 15]^. Besides, some bacterial structures, such as flagellum and pilus, can also serve as phage receptors ^[16, 17]^. The carbohydrate and lipid are widely distributed on the host cell surfaces and are easily taken as receptors by viruses ^[18]^. Compared to these molecules, proteins are more suitable as viral receptors due to stronger affinity and higher specificity for viral attachment ^[19]^. The viral protein receptors are highly selective. Among over 2,000 human plasma membrane proteins, less than 100 proteins have been reported to be viral receptors ^[20]^. Previous studies have shown that the proteins abundant on the host cell surface, or with relatively low affinity for their natural ligands, or having high N-glycosylation and large number of interaction partners in the PPI network are preferred by viruses as receptors ^[18, 20, 21]^. However, most research about characterizing viral protein receptors has been done on mammalian viruses, and little is known about the molecular characteristics of protein receptors of phages.

Here, we systematically analyzed the phage-host receptor interactions and characterized the phage protein receptors (PPRs) by manually curating an up-to-date and high-quality database containing 355 pairs of phage-host receptor interactions. This study helps to understand the mechanism underlying phage–receptor interactions, as well as predict and identify phage receptors.

## Materials and methods

### Database of phage-host receptor interactions

The data of phage-host receptor interactions were compiled from two sources: firstly, 109 pairs of phage-receptor interactions were directly obtained from the Phage Receptor Database ^[13]^ on May 1^st^, 2019; secondly, the literatures related to phage receptors were downloaded from NCBI PubMed database by searching “phage receptor” [TIAB] or “bacteriophage receptor”[TIAB] on June 4^th^, 2019. 246 pairs of phage-receptor interactions were manually extracted from the literatures. In total, 355 pairs of phage-host receptor interactions were collected for further analysis. The details of the phage-receptor interactions are public available at the database of phageReceptor (http://www.computationalbiology.cn/viralRecepetor/index.html).

### Escherichia coli proteome and location of proteins

The reference proteome of *Escherichia coli* (*E. coli*) str. K-12 substr. MG1655 (Proteome ID: UP000000625) was downloaded from the database of UniProt Proteome ^[22]^ on May 16, 2019. The location of proteins in *E. coli* and other species was inferred based on the field of “Subcellular location” of proteins provided in the database of UniProtKB, or on the Gene Ontology (GO) Cellular Component annotations of proteins. In GO annotations, the proteins annotated with GO terms containing the case-insensitive words of “membrane” were considered to be located on the membrane; those annotated with GO terms containing the case-insensitive words of “cell outer membrane” were considered to be located on the outer membrane; those annotated with GO terms containing the case-insensitive words of “cell inner membrane”, “cell membrane” or “plasma membrane” were considered to be located on the plasma membrane; and those annotated with GO terms of “bacterial-type flagellum hook” were considered to be located in the flagellum.

### Analysis of structural features of PPRs

The transmembrane structures of the PPRs were derived from the field of “Transmembrane” of proteins in the database of UniProtKB. Besides, the TMHMM Server (v.2.0) ^[23]^ was used to predict the transmembrane helix of the PPRs. The Pfam domains included in the PPRs were derived from the field of “Pfam” of proteins in the database of UniProtKB. Besides, InterProScan (version 5) ^[24]^ was used to identify Pfam domains in PPRs.

### Functional enrichment analysis

The GO function and KEGG pathway enrichment analysis for the *E. coli* PPRs were conducted with the functions of enrichGO() and enrichKEGG() in the package “clusterProfiler” (version 3.10.0) ^[25]^ in R (version 3.5.1).

### PPI network analysis

The *E. coli* PPI network (PPIN) was built based on the PPIs of the *E. coli* str. K-12 substr. MG1655 (NCBI taxon-Id: 511145) downloaded from the STRING database ^[26]^ on May 16, 2019. Only the PPIs with median confidence (combined score ≥ 400) were kept for analysis. The PPIN included 175,845 PPIs, and 4,121 proteins which could be mapped to UniProt identifiers. The number of membrane proteins, plasma membrane proteins, outer membrane proteins and phage receptor proteins included in the PPIN was 1519, 1183, 120 and 15, respectively. Two measures were used to analyze the importance of proteins in the *E. coli* PPIN. The first is the node degree, which is defined as the number of connections the node has to other nodes in the PPIN; the other is the node betweenness, which is defined as the number of shortest paths that pass through the node. The degree and betweenness of each protein in the PPIN were calculated with functions of degree() and betweenness() in the package “igraph” (version 1.2.2) ^[27]^ in R (version 3.5.1). The network was displayed with Cytoscape (version 3.7.0) ^[28]^.

### Analysis of the expression level of PPRs and other genes in E. coli

The expression level of PPRs and other genes in *E. coli* were obtained from LaCroix’s work ^[29]^ (GEO accession number: GSE61327), during which the expressions of genes of wild-type *E. coli* str. K-12 substr. MG1655 (the strain used for PPI analysis) under normal conditions were measured by RNA-seq. The average expression level of each gene in two replicates was used. The gene expression level was measured in reads per kilobase per million mapped reads (RPKM).

### Statistical analysis

All the statistical analysis was conducted in R (version 3.5.1) ^[30]^. The wilcoxon rank-sum test was conducted with the function of wilcox.test(). P-values < 0.05 were considered to be statistically significant.

## Results

### Summary of phage-host receptor interactions

To characterize the phage receptors, we manually curated a high-quality database of 355 pairs of phage-host receptor interactions (details in Materials and methods), including 280 unique viral species or sub-species from 11 viral families and 64 bacterial species. 88.9% of the viruses included in the database were double-stranded DNA viruses, while the remaining viruses were single-stranded DNA viruses (7.1%), double-stranded RNA viruses (1.4%), single-stranded RNA viruses (1.4%) and unclassified viruses (1.2%). Viruses belonging to the families of Siphoviridae, Podoviridae and Myoviridae were most abundant and accounted for 84% of all viruses in the database (Figure S1). The bacteria included 48 Gram-negative bacteria and 16 Gram-positive bacteria, and *E. coli* was the most frequently observed bacterial species in the database which took part in 28% of phage-receptor interactions.

Multiple types of receptors were observed for phages, including proteins, sugar, acid and bacterial structures such as flagellum, pilus and exopolysaccharide capsular. Sugars were most frequently used by phages as receptors (49%), followed by proteins (33%), bacterial structures (14%) and acid (4%). Most viruses (92%) used only one type of receptors. The receptor usage was further analyzed for three largest viral families, i.e., Siphoviridae, Podoviridae and Myoviridae. Viruses in these families had similar preferences towards receptors, with sugar used mostly widely (Figure 1A). Interestingly, viruses infecting Gram-negative and Gram-positive bacteria had different preferences towards receptors. For example, in the family of Siphoviridae, most viruses infecting Gram-negative bacteria used proteins (blue) as receptors, and none of them used acid (cyan) as receptors; while most viruses infecting the Gram-positive bacteria used sugars (red) as receptors, and none of them used bacterial structures (yellow) as receptors.

**Figure 1.**
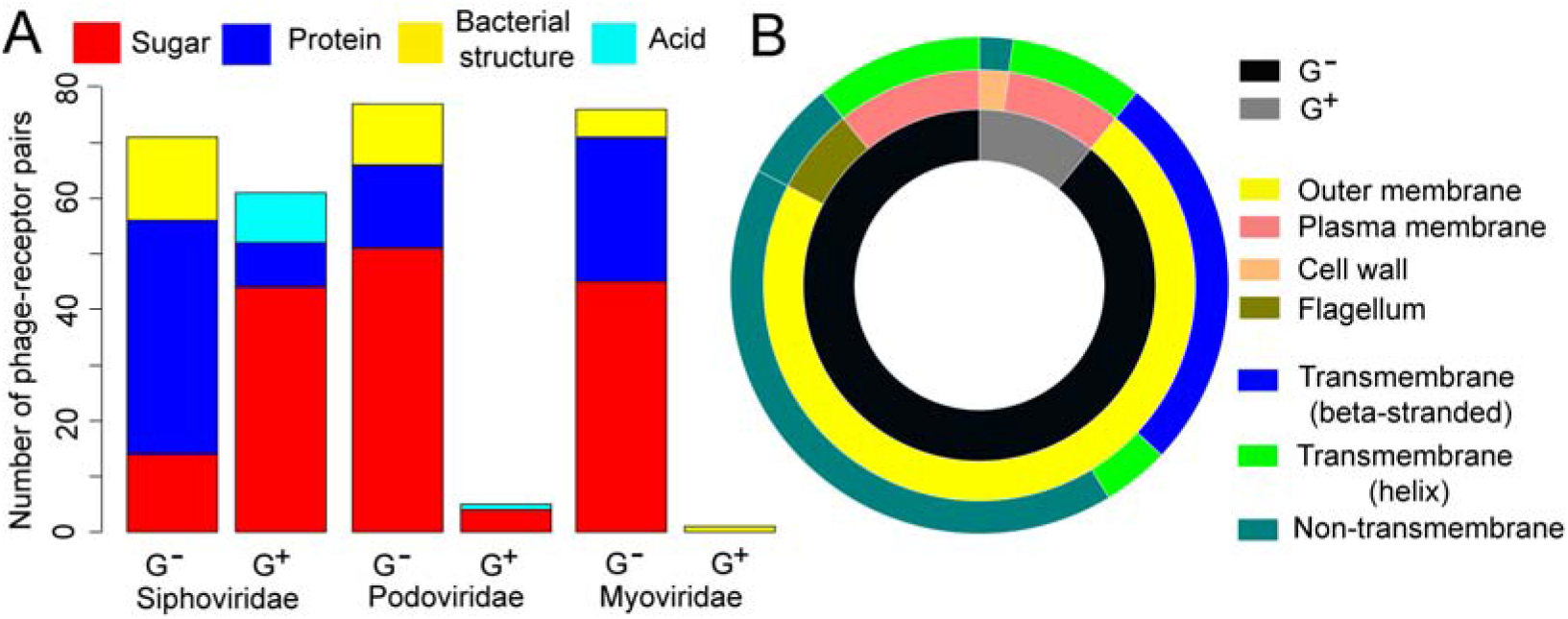
The receptor usage in three largest viral families (A) and the location of PPRs (B). A total of 48 protein receptors were identified for the phages (Table S1). They were distributed in 15 bacterial species, among which *E. coli* had the largest number of protein receptors (18/48) (Figure S2). Analysis of the association between phages and their protein receptors showed 84% of phages used one specific receptor only (Figure S3). Next, the protein receptor usage on the level of viral family was analyzed. For three largest viral families in the database, the viruses in each of them used two or more sets of protein receptors (Figure S3), suggesting that different viruses of the same family tend to use different receptors. For example, in the family of Siphoviridae, a total of 22 protein receptors were used. On the contrary, some viruses of different families shared the same protein receptor. For example, Phage 434 and Escherichia virus T4, from the family of Siphoviridae and Myoviridae respectively, both took the outer membrane porin C (ompC) (marked by a star in Figure S3) as the receptor in infecting the *E. coli.* Twenty protein receptors were used by more than one virus (Figure S3). Especially, some proteins, such as ompC and ferrichrome outer membrane transporter in *E. coli,* were used by more than five phages as receptors.

### Location of PPRs

We continued our study to analyze the location of PPRs (Figure 1B). Among PPRs in the Gram-negative bacteria, 80% were located on the outer membrane, among which 12 receptors had a transmembrane beta barrel and 2 receptors had a transmembrane helix, and the remaining 20% were located on the plasma membrane or flagellum. As for five PPRs in the Gram-positive bacteria, four of them were located in the plasma membrane, each of which had six transmembrane helixes, and the rest was located in the S-layer, which is a paracrystalline protein thin layer attached to the outermost portion of the cell wall.

### PPRs were diverse in structure

Next, we investigated the structural characteristics of PPRs by analyzing the protein domains. The PPRs consisted of 69 domains based on the Pfam database (Table S2), which can be categorized into 33 Pfam families. Especially, Gram-negative porin, TonB-dependent Receptor and TonB-dependent Receptor Plug Domain were observed in more than five PPRs. Comparison of the Pfam domains included in the PPRs and mammalian virus protein receptors showed no overlap. Besides, little sequence homology was observed between PPRs and mammalian virus protein receptors (data not shown).

### Functional enrichment analysis of PPRs in *E. coli*

We next characterized the functions of PPRs in *E. coli,* since *E. coli* is the most characterized bacterial species, and as mentioned above, *E. coli* had most PPRs in our database (18/48) while 12 of the remaining 30 PPRs in other species were homologs of PPRs in *E. coli* (Table S3). The function enrichment analysis was conducted based on the databases of GO and Kyoto Encyclopedia of Genes and Genomes (KEGG). According to the results of GO analysis, in the domain of Cellular Component, the PPRs in *E. coli* were mainly enriched in the outer membrane and envelope; in the domain of Biological Process, the PPRs were mainly enriched in GO terms related to transport, localization and virus’s entry into host cells; in the domain of Molecular Function, the PPRs were mainly enriched in GO terms related to binding, channel and transporter activity. For the KEGG pathway, only the beta-Lactam resistance and Two-component system pathways were significantly enriched in the PPRs in *E. coli*.

**Figure 2.**
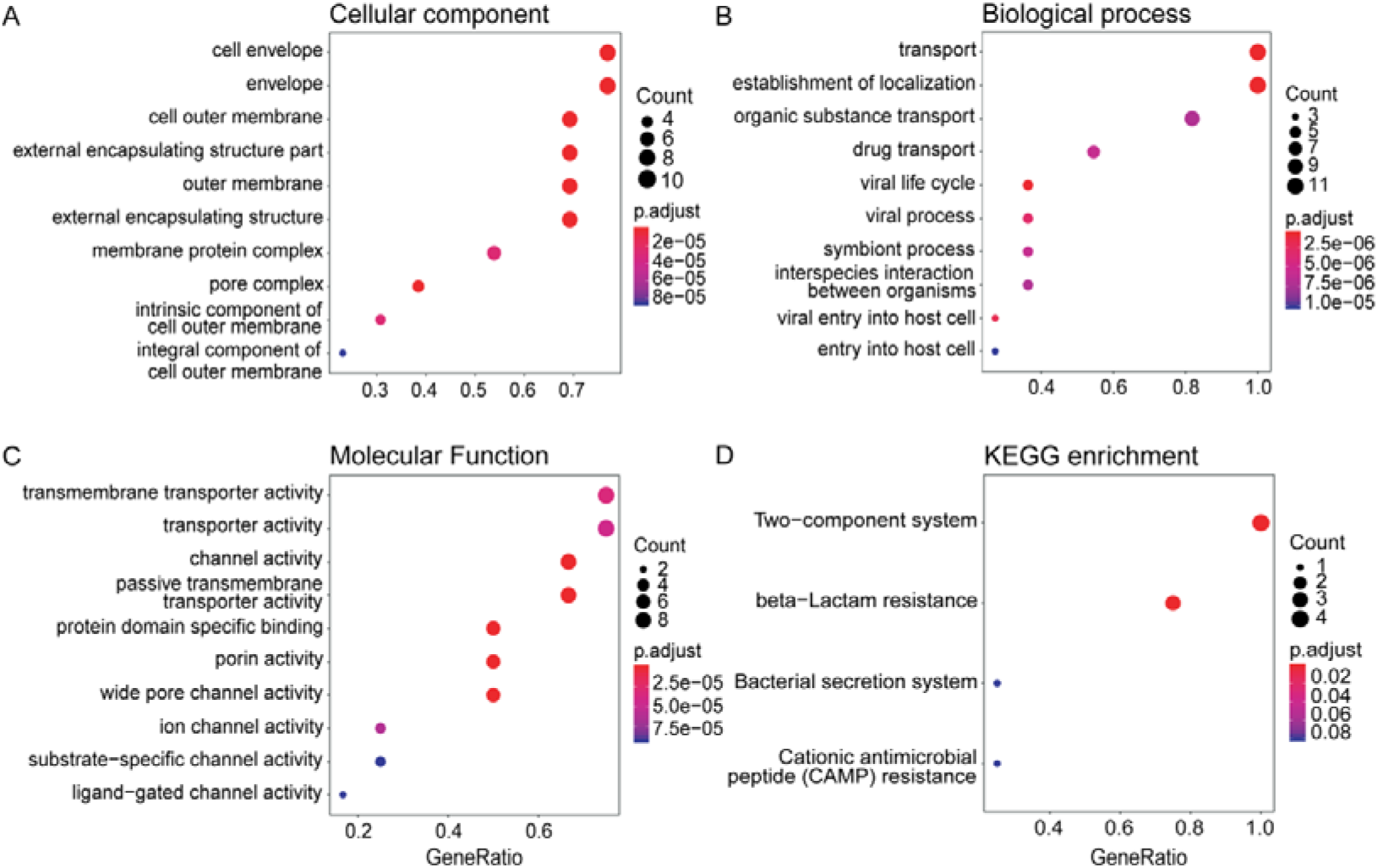
The GO and KEGG pathway enrichment analysis of PPRs in *E. coli.* Only the top 10 enriched GO terms were shown.

### PPI analysis of PPRs in *E. coli*

Then, the role of PPRs in the *E. coli* PPIN was investigated. Firstly, an *E. coli* PPIN was constructed which included 4,121 E. *coli* proteins and 175,845 interactions (Materials and methods). To evaluate the role of proteins in the *E. coli* PPIN, the degree of each protein was calculated, which is defined as the number of connections the protein has to other proteins in the *E. coli* PPIN. Because the PPRs in *E. coli* were exclusively located on the outer membrane (16/18) and the plasma membrane (2/18), the degrees of PPRs were compared to those of other outer membrane and plasma membrane proteins. As was shown in Figure 3A, the degrees of outer membrane proteins were slightly higher than those of plasma membrane proteins, membrane proteins, and whole proteins in *E. coli*. While the PPRs, which were part of outer membrane and plasma membrane proteins, had a median of 124 interaction partners in the PPIN (Figure 3A), which was 2 times that of outer membrane proteins, plasma membrane proteins, membrane proteins and all proteins in *E. coli*.

**Figure 3.**
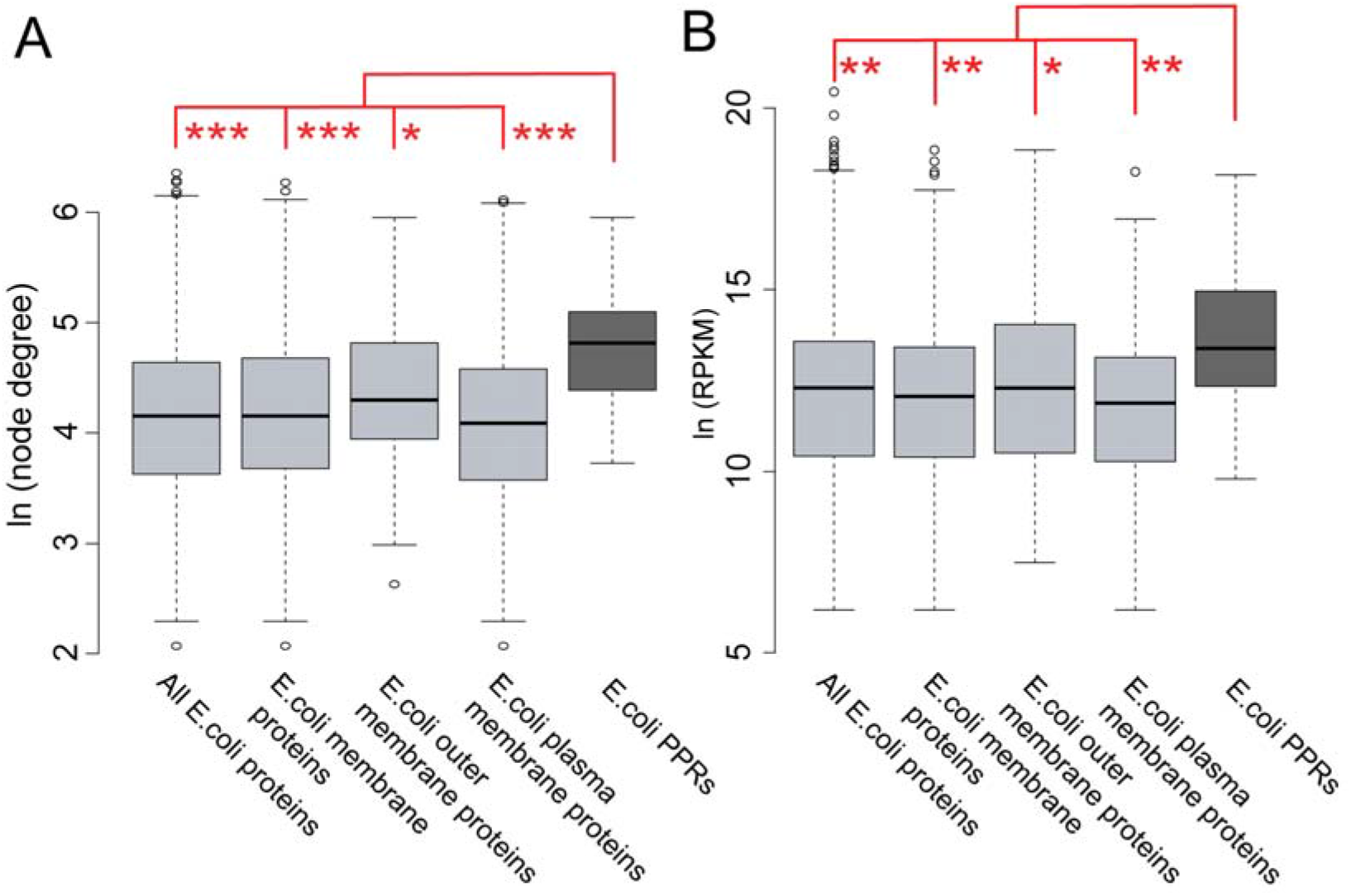
Distribution of the node degrees of PPRs in the *E. coli* PPIN (A) and the expression levels of PPRs in *E. coli* (B), and its comparison to other sets of proteins, including *E. coli* outer membrane proteins, plasma membrane proteins, membrane proteins and all *E. coli* proteins. The expression level was measured in RPKM. Both the node degree and expression level were transformed in natural logarithm for clarity. “***”, “**”, “*” refer to p-values of smaller than 0.001, 0.01, 0.05, respectively, in the Wilcoxon rank-sum test.

For robustness of the results, the node betweenness (another measure of node centrality in network) of each protein was also calculated based on the E. *coli* PPIN. The results showed the betweennesses of E. coli PPRs were significantly higher than those of other sets of proteins (Figure S4), further demonstrating the importance of PPRs in *E. coli* PPIN.

### High expression of PPRs in *E. coli*

As viruses compete with host proteins in binding to receptors, proteins with relatively high expressions are supposed to be preferred by phages as receptors. Therefore, the expressions of *E. coli* PPRs, and those of outer membrane proteins, plasma membrane proteins, membrane proteins and all proteins in *E. coli* were collected and compared with each other (Materials and methods). As shown in Figure 3B, the expressions of outer membrane proteins were slightly larger than those of plasma membrane proteins, membrane proteins and all proteins in *E. coli.* As expected, the expression level of PPRs was nearly 100% higher than that of outer membrane proteins, with a median of 675,299.5 RPKM (13.4 in transformed natural logarithm).

## Discussion

This study systematically analyzed the associations between phage and host receptors. Multiple types of receptors were identified, including sugars, proteins, acid and bacterial structures. Sugars and proteins are widely used by phages as receptors. Interestingly, the receptor usage of phages in Gram-positive bacteria is different from that in Gram-negative bacteria, for which a potential explanation is due to different cell-wall structures of the two types of bacteria ^[31]^. For example, no viruses take acid as receptors when infecting the Gram-negative bacteria, because there is no acid on the cell wall of the Gram-negative bacteria; viruses infecting the Gram-positive bacteria use a smaller ratio of proteins as receptors than those infecting the Gram-negative bacteria, because there is few proteins in the cell wall of the Gram-positive bacteria.

The PPRs were systematically characterized by location, structure, function, protein-protein interaction and gene expression. Similar to the mammalian virus protein receptors, the PPRs were structurally diverse. They had no common protein domains with mammalian virus protein receptors. For the functional enrichment analysis, besides the molecular function of binding and biological process of viral entry into host cell which were also enriched in the human virus protein receptors ^[20]^, the molecular functions of channel and transporter activity, and the biological processes of transport and location, were enriched in PPRs in *E. coli*. This suggests PPRs may be involved in transportation of viral genetic materials into host cells.

The PPRs in *E. coli* have much larger node degrees and betweennesses than other *E. coli* proteins. This indicates the important roles of viral receptors in the host cells. In addition, the expression level of *E. coli* RPRs is higher than other proteins, which can facilitate their interactions with more proteins. These features are similar to those observed for the human virus protein receptors which also have multiple interaction partners in the human PPIN and are highly expressed in common human tissues^[20]^. Considering the huge difference between bacteria and mammals, and the diversity of phages and mammalian viruses used in this study and previous studies ^[20]^, centrality in the host PPIN may be a common feature of viral host receptors. Since viruses have to compete with host proteins for binding to the receptors, the proteins with less interaction partners are supposed to be more suitable receptors for viruses. Previously we hypothesized the virus receptor proteins are closely related to the “door” of the host cell, so that viruses have to interact with them for entry into the host cell ^[20]^. This hypothesis is supported by the functional analysis of the PPRs in *E. coli*, as they have significantly enriched functions of channel and transporter activity. Actually, 13 of 18 PPRs in *E. coli* belong to transporter, channel or porin proteins, which may facilitate viral entry.

Phage therapy, i.e., the therapeutic use of phages to treat pathogenic bacterial infections, has been considered as a promising strategy for treatment of drug-resistant bacterial infections in human medicine, veterinary science, and agriculture ^[32]^. Better understanding of the phage-host receptor interactions would promote the development and application of phage therapy. This study shows that phages can take multiple types of molecules as receptors. Genetic modification of phages, such as recombination of receptor-binding protein domains from multiple phages ^[12, 33]^, may help them recognize novel types of receptors and broaden the host range. Besides, most PPRs in Gram-negative bacteria are located on the outer membrane. Considering the small number of outer membrane proteins, the ratio of receptor proteins is relatively high in the outer membrane proteins, which suggests phages tend to take outer membrane proteins as receptors when infecting Gram-negative bacteria. This can help identify PPRs in the drug-resistant bacteria.

Nevertheless, this study has several limitations. Firstly, the database of phage-host receptor interactions, especially the PPRs, was still limited in its size due to the difficulties in identifying phage receptors. Fortunately, several common features were identified for the PPRs, such as the structure diversity and centrality in PPIN. These features may help identify novel PPRs. Secondly, the phage-host receptor interactions and PPRs used in this study were biased towards the Gram-negative bacteria. Thus, more efforts are needed to identify the interactions between phages and host receptors in Gram-positive bacteria.

In conclusion, this study systematically analyzed the phage-host receptor interactions and characterized the PPRs. The PPRs were found to be structurally diverse, play a central role in PPIN and are highly expressed, which are similar to mammalian virus protein receptors. These features could help development of effective methods for phage receptor identification. This study can not only help understand the mechanisms underlying virus-host interactions, but also help for development and application of phage therapy.

## ACKNOWLEDGEMENTS

This work was supported by the National Key Plan for Scientific Research and Development of China (2016YFD0500300), Hunan Provincial Natural Science Foundation of China (2018JJ3039 and 2019JJ50035), the National Natural Science Foundation of China (31671371) and the Chinese Academy of Medical Sciences (2016-I2M-1-005), and the Fundamental Research Funds for the Central Universities of China (531107051162).

The authors have declared that no competing interests exist.

## References

[1] Letarov AV, Kulikov EE. Adsorption of bacteriophages on bacterial cells. Biochemistry-Moscow+, 2017, 82: 1632–1658

[2] Garcia-Doval C, van Raaij MJ. Bacteriophage receptor recognition and nucleic acid transfer. Sub-cellular biochemistry, 2013, 68: 489–518

[3] Grath SM, Sinderen Dv. Bacteriophage: Genetics and molecular biology. Caister Academic Press, 2007

[4] Gonzalez F, Helm RF, Broadway KM, et al. More than rotating flagella: Lipopolysaccharide as a secondary receptor for flagellotropic phage 7-7-1. Journal of bacteriology, 2018, 200:

[5] Gregory AC, Zayed AA, Conceicao-Neto N, et al. Marine DNA viral macro- and microdiversity from pole to pole. Cell, 2019, 177: 1109-+

[6] Paez-Espino D, Eloe-Fadrosh EA, Pavlopoulos GA, et al. Uncovering earth’s virome. Nature, 2016, 536: 425-+

[7] Spinelli S, Veesler D, Bebeacua C, et al. Structures and host-adhesion mechanisms of lactococcal siphophages. Front Microbiol, 2014, 5:

[8] Doss J, Culbertson K, Hahn D, et al. A review of phage therapy against bacterial pathogens of aquatic and terrestrial organisms. Viruses-Basel, 2017, 9:

[9] Torres-Barcelo C, Hochberg ME. Evolutionary rationale for phages as complements of antibiotics. Trends in microbiology, 2016, 24: 249–256

[10] Dalmasso M, Hill C, Ross RP. Exploiting gut bacteriophages for human health. Trends in microbiology, 2014, 22: 399–405

[11] Barr JJ. Precision engineers: Bacteriophages modulate the gut microbiome and metabolome. Cell host & microbe, 2019, 25: 771–773

[12] de Jonge PA, Nobrega FL, Brouns SJJ, et al. Molecular and evolutionary determinants of bacteriophage host range. Trends in microbiology, 2019, 27: 51–63

[13] Silva JB, Storms Z, Sauvageau D. Host receptors for bacteriophage adsorption. Fems Microbiol Lett, 2016, 363:

[14] Casasnovas JM. Virus-receptor interactions and receptor-mediated virus entry into host cells. 2013[

[15] Mazzon M, Mercer J. Lipid interactions during virus entry and infection. Cellular microbiology, 2014, 16: 1493–1502

[16] Kim S, Rahman M, Seol SY, et al. Pseudomonas aeruginosa bacteriophage pa1empty set requires type iv pili for infection and shows broad bactericidal and biofilm removal activities. Appl Environ Microb, 2012, 78: 6380–6385

[17] Attridge SR, Fazeli A, Manning PA, et al. Isolation and characterization of bacteriophage-resistant mutants of vibrio cholerae o139. Microb Pathogenesis, 2001, 30: 237–246

[18] Dimitrov DS. Virus entry: Molecular mechanisms and biomedical applications. Nat Rev Microbiol, 2004, 2: 109–122

[19] Baranowski E, Ruiz-Jarabo CM, Domingo E. Evolution of cell recognition by viruses. Science, 2001, 292: 1102–1105

[20] Zhang Z, Zhu ZZ, Chen WJ, et al. Cell membrane proteins with high n-glycosylation, high expression and multiple interaction partners are preferred by mammalian viruses as receptors. Bioinformatics, 2019, 35: 723–728

[21] Wang JH. Protein recognition by cell surface receptors: Physiological receptors versus virus interactions. Trends in biochemical sciences, 2002, 27: 122–126

[22] Bateman A, Martin MJ, O’Donovan C, et al. Uniprot: The universal protein knowledgebase. Nucleic Acids Res, 2017, 45: D158–D169

[23] Krogh A, Larsson B, von Heijne G, et al. Predicting transmembrane protein topology with a hidden markov model: Application to complete genomes. Journal of molecular biology, 2001, 305: 567–580

[24] Quevillon E, Silventoinen V, Pillai S, et al. Interproscan: Protein domains identifier. Nucleic Acids Res, 2005, 33: W116–120

[25] Yu GC, Wang LG, Han YY, et al. Clusterprofiler: An r package for comparing biological themes among gene clusters. Omics, 2012, 16: 284–287

[26] Szklarczyk D, Franceschini A, Wyder S, et al. String v10: Protein-protein interaction networks, integrated over the tree of life. Nucleic Acids Res, 2015, 43: D447–D452

[27] Csardi G, Nepusz T. The igraph software package for complex network research, interjournal, complex systems 1695. 2006, http://igraph.org

[28] Shannon P, Markiel A, Ozier O, et al. Cytoscape: A software environment for integrated models of biomolecular interaction networks. Genome research, 2003, 13: 2498–2504

[29] LaCroix RA, Sandberg TE, O’Brien EJ, et al. Use of adaptive laboratory evolution to discover key mutations enabling rapid growth of escherichia coli k-12 mg1655 on glucose minimal medium. Appl Environ Microb, 2015, 81: 17–30

[30] Team RC. R: A language and environment for statistical computing. R foundation for statistical computing, vienna, austria. 2019, https://www.R-project.org/

[31] Johnson JW, Fisher JF, Mobashery S. Bacterial cell-wall recycling. Ann Ny Acad Sci, 2013, 1277: 54–75

[32] Cisek AA, Dabrowska I, Gregorczyk KP, et al. Phage therapy in bacterial infections treatment: One hundred years after the discovery of bacteriophages. Curr Microbiol, 2017, 74: 277–283

[33] Ando H, Lemire S, Pires DP, et al. Engineering modular viral scaffolds for targeted bacterial population editing. Cell Syst, 2015, 1: 187–196

